# A Graph Convolutional Network-based screening strategy for rapid identification of SARS-CoV-2 cell-entry inhibitors

**DOI:** 10.1101/2021.12.08.471787

**Authors:** Peng Gao, Miao Xu, Qi Zhang, Catherine Z Chen, Hui Guo, Yihong Ye, Wei Zheng, Min Shen

## Abstract

The cell entry of SARS-CoV-2 has emerged as an attractive drug development target. We previously reported that the entry of SARS-CoV-2 depends on the cell surface heparan sulfate proteoglycan (HSPG) and the cortex actin, which can be targeted by therapeutic agents identified by conventional drug repurposing screens. However, this drug identification strategy requires laborious library screening, which is time-consuming and often limited number of compounds can be screened. As an alternative approach, we developed and trained a graph convolutional network (GCN)-based classification model using information extracted from experimentally identified HSPG and actin inhibitors. This method allowed us to virtually screen 170,000 compounds, resulting in ∼2000 potential hits. A hit confirmation assay with the uptake of a fluorescently labeled HSPG cargo further shortlisted 256 active compounds. Among them, 16 compounds had modest to strong inhibitory activities against the entry of SARS-CoV-2 pseudotyped particles into Vero E6 cells. These results establish a GCN-based virtual screen workflow for rapid identification of new small molecule inhibitors against validated drug targets.

**Graphical TOC Entry:** 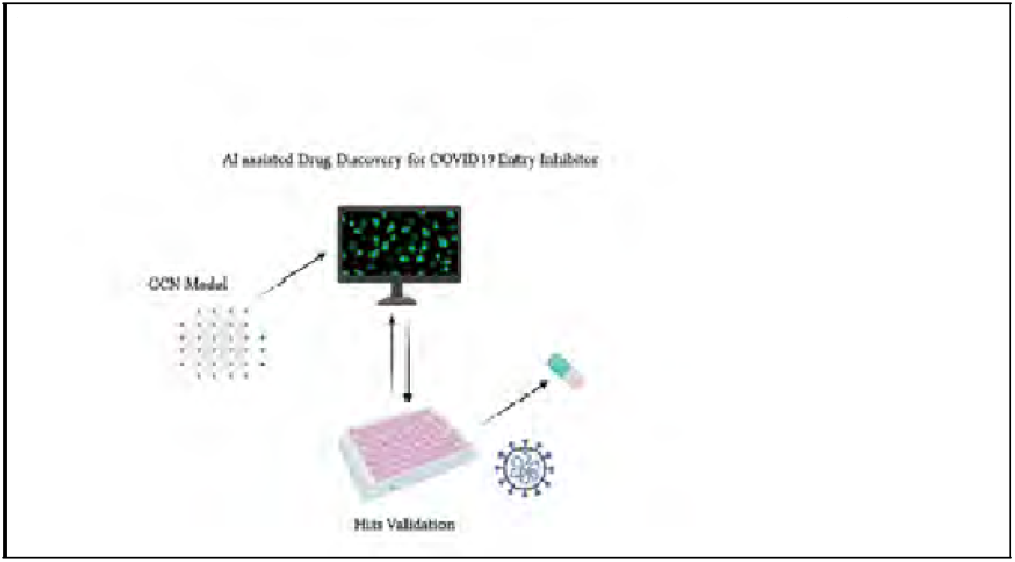

## Introduction

Since the outbreak of the COVID19 pandemic, global communities have suffered a significant loss of lives and economic growth. Although the development of COVID vaccines can significantly contain the spreading of SARS-CoV-2, the virus is constantly evolving into more infectious and transmissible variants (e.g., the delta strain), resulting in infrequent breakthrough infections among vaccinated people.^1–5^ The constant increase of hospitalized patients in the USA and around the world despite the rollout of the vaccination programs has summoned the need to develop potent small molecule therapeutics for COVID patients.

The cellular entry of SARS-CoV-2 is one of the key steps in the viral life cycle that represents a hot target for small molecule inhibitors.^6,7^ The entry of SARS-CoV-2 requires the interaction of the glycosylated viral Spike protein with the angiotensin-converting enzyme 2 (ACE2) receptor on the cell surface. ^8–12^ Previously, we and others identified the cell surface heparan sulfate proteoglycans (HSPGs) as a critical factor that facilitate the entry of SARS-CoV-2 virions. 8,9 We further showed that HSPGs also facilitate the uptake of other positive charge-bearing endocytic cargos such as supercharged GFP and preformed *α*-Synuclein pathogenic fibrils.^13^ HSPGs are a family of glycoproteins bearing one or more negatively charged polysaccharide chains consisting of repeated heparan sulfate disaccharide units. Most HSPG family members are anchored to the cell surface either as a single spanning membrane protein (e.g., Syndecans) or Glycosylphosphatidylinositol (GPI) -anchored protein (e.g., Glypicans). Due to the enrichment of negatively charged sulfate groups, HSPGs can effectively serve as an attachment anchor to increase the surface dwell time for endocytic cargos bearing positive charges, facilitating their engagement with a downstream receptor.^6,13,14^The internalization of HSPG cargos also requires the cortex actin network, which maintains plasma membrane dynamics to promote the maturation of clathrin-coated pits.^13^

We recently conducted a drug repurposing screen, which identified 8 drugs that inhibited HSPG-dependent entry of SARS-CoV-2 virions. Intriguingly, despite structural dissimilarity, several of the identified drugs can all bind directly to heparin, a heparan sulfate analog, suggesting that they may target the polysaccharide chain on the cell surface of HSPG to inhibit viral entry. In addition to heparin-binding drugs, two structurally unrelated drugs, Sunitinib and BNTX, can both effectively disrupt the actin filaments underlying the plasma membrane (cortex actin) to inhibit HSPG-mediated endocytosis. ^9–12^

While drug repurposing screen is an effective strategy to rapidly adopt existing drugs for new therapeutic uses, the original target(s) of the approved drugs often reduces their therapeutic specificity, which may cause undesired side effects for treating diseases like viral infection. For example, as a heparan sulfate binding compound, mitoxantrone delivers the most potent antiviral activity in vitro. However, because mitoxantrone was originally approved as anti-cancer chemotherapy via targeting the DNA topoisomerase,^15^ cytotoxicity associated with DNA replication inhibition is an obvious concern.

We postulate that drugs bearing partial structural elements from the identified HSPG and actin inhibitors may retain the endocytosis inhibition function but fail to act on the original target(s), and therefore be more specific. In this regard, conventional structure-activity-relationship (SAR) studies, albeit labor-intensive and time-consuming, often yield unpredictable results. To identify additional inhibitors targeting HSPG-mediated viral entry, we developed a graph convolutional network (GCN)-based classification approach. GCN can efficiently translate 3D structures into molecular graphs composed of nodes and edges, and then utilize these graphs to extract spatial information to achieve accurate molecular classification and properties predictions. ^16–19^ Compared to other traditional computational methods based on molecular dynamics (MD) simulations or density functional theory (DFT), the computational cost of GCN is substantially lower. These features allowed us to rapidly screen 17,000 compounds in several NCATS libraries. From these libraries, we identified and confirmed a set of compounds (256) as inhibitors of HSPG-dependent endocytosis with the most potent IC50 value at 0.95 µM. Further testing with a SARS-CoV-2 pseudotyped particle entry assay confirmed 16 compounds as entry inhibitors.

## Methods

### Computational details

#### GCN model

GCN-based approaches display considerable robustness for structural elucidations, ^16–19^ because it could fully utilize the molecular graphs for information extraction with substantially reduced computational cost. ^20–27^ In addition, such an architecture is also flexible enough to include different chemical knowledge for specific assignments.^20,28–37^ In this study, we employed the self-developed GCN package for activity classifications. The workflow of the applied GCN was described in Figure 1. For any given drug molecule, its structural information was contained in the simplified molecular-input line-entry system (SMILES) string, and GCN can transform the molecular graph into a set of numerical descriptors for computational processing.

**Figure 1:**
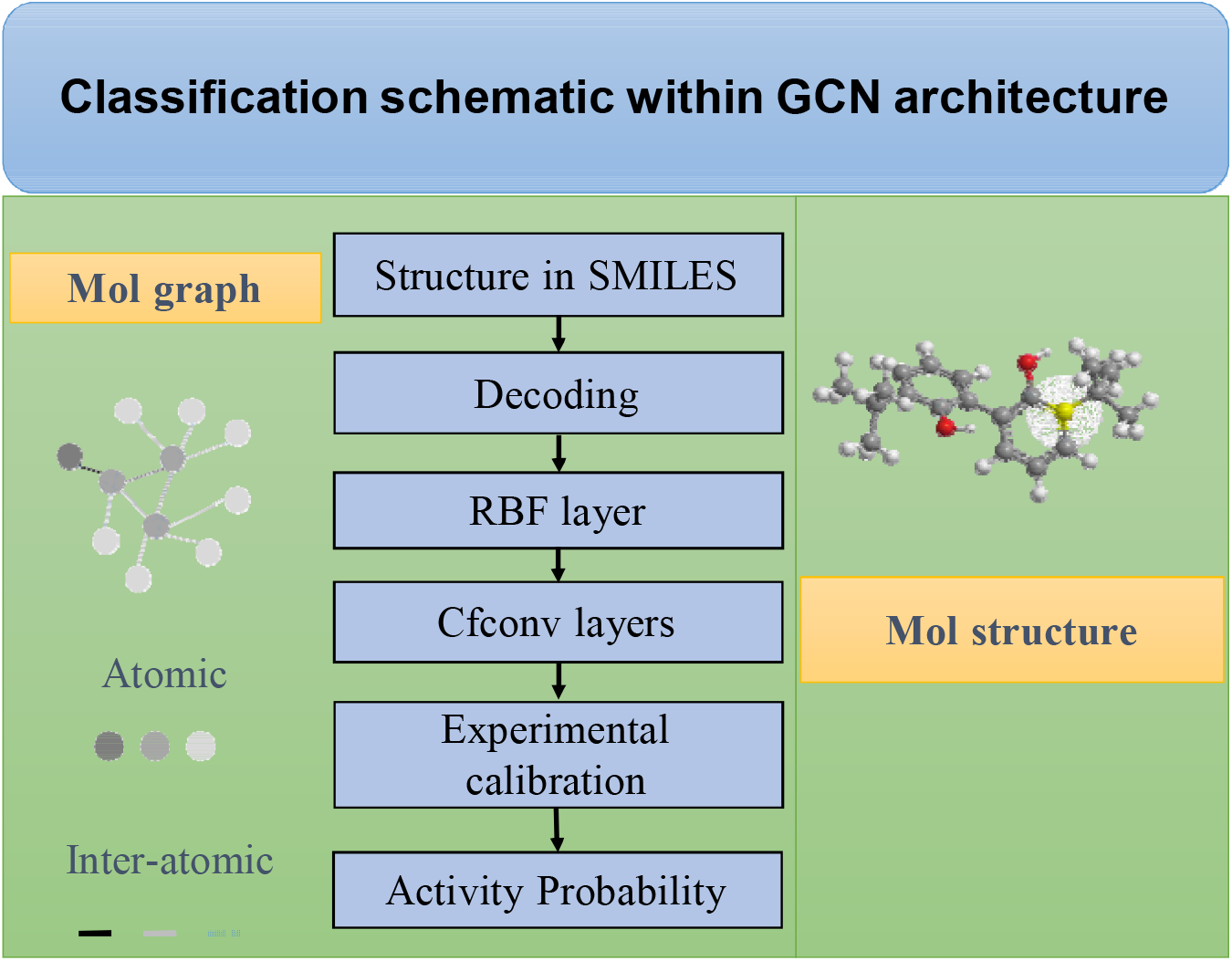
The architecture of GCN classification model for virtual screenings.

All the collected SMILES strings of drug molecules were first translated into molecular graphs through the *TencentAlchemyDataset* within Deep Graph Library (DGL) library. ^38,39^ Each drug molecule is composed of edges and nodes within 3D space. Within the framework of GCN, the nodes are more associated with atomic features, while the edges are corresponding to bonding descriptors. Thus, molecular graphs with full connections can reasonably represent drugs’ 3D structures. And with the numerically solved drugs’ structures, related molecular properties can be well mapped. In fact, within any molecular or fragmentary graphs, all the connections between every two atoms are fully utilized for information extraction; the specific values were recorded in distance tensors at the radial basis function (RBF) layer, guaranteeing there is no omission of important structural information. In addition, within GCN model, to decently solve molecular graphs at atomic level, multiple continuous-filter convolutions (cfconv) layers were employed to optimize and record the inter-atomic evolution. For instance, at ***k***+***1*** layer, the ***i*** th atom’s evolution can be expressed with the following equation:

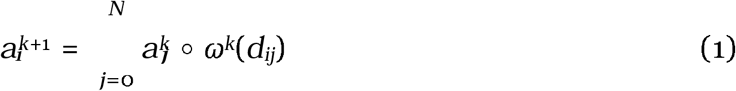

in which, ∘ represents element-wise multiplication, and *ω*^*k*^ is the filter-generation that can map the atoms’ descriptions to the filter bank. To efficiently control the evolution accuracy via the applied the filter values, a Gaussian-type function, *gauss*_*k*_, was employed, which can be expressed with the following equation:

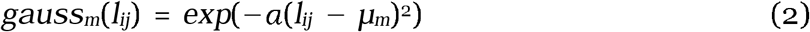

where, *µ*_*m*_ is the pre-set value of cutoff, and *l*_*ij*_ represents the bonding distance among the ***i*** th atom and ***j*** th atom. The *a* is attributed to hyper parameters, and it was set to 0.1 in this study. ^40^

For any predictive property or classification task, the computed value, *Pro*, by GCN model is calibrated with respect to experimental measurement, *Pro*, and the accuracy can be well indicated by the squared loss function, as shown below:

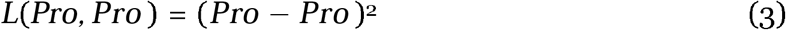

In this study, we applied the developed GCN package for drugs activity classification; however, it is worth noting that this promising architecture is also able to include various kinds of chemical & physical knowledge for more challenging structural assignments.

#### Data set

We applied the above-described GCN model to a previously reported COVID-19 related drug screening, which identified drugs that block HSPG-dependent entry of *α*-Synuclein fibrils. Classification algorithm was based on NCATS’ collected activity values. The model was first trained by the collected data, which consisted of 3,832 compounds. Among them, 367 compounds show activities and 3,465 are inactive. These compounds were randomly divided with a ratio of 9:1; and 90% was used as the training set, and the remaining 10% as the test set. The trained GCN model was validated by the compounds in the test set, which scored an accuracy of 99.5%. The trained model was then used to screen more than 170,000 compounds contained in three independent libraries, Genesis, Sytravon, and NPACT, none of which had been experimentally screened by endocytosis or SARS-CoV-2 PP entry assays.

### *a*-Synuclein fibrils uptake assay and drug verification

Fluorescence labeled alpha-synuclein fibrils were generated as previously described. ^13^ HEK293T cells were dispensed into black, clear-bottom 1536-well microplates (Greiner BioOne, # 789092-F)) at 5000cells/well in 5L media with 200nM pHrodo red-labeled *α*-Syn fibrils and incubated at 37°C, 5% CO2, 85% humidity overnight (∼16 h). Compounds picked from the virtual screen were titrated 1:3 with 11 points in DMSO and transferred to assay plates at a volume of 23 nl/well by an automated pintool workstation (Wako Automation, San Diego, CA). After 24 h of incubation, the fluorescence intensity of pHrodo red was measured by a CLARIOstar Plus plate reader (BMG Labtech). Data was normalized using the wells with cells containing 200nMpHrodo red-labeled Syn fibrils as 100% and the wells without cells as 0%.

#### Image processing and statistical analyses

Confocal images were processed using the Zeiss Zen software. To measure fluorescence intensity, we used the Fiji software. Images were converted to individual channels and regions of interest were drawn for measurement. Statistical analyses were performed using either Excel or GraphPad Prism 9. Data are presented as means ± SEM, which was calculated by GraphPad Prism 9. P values were calculated by Student’s t-test using Excel. Nonlinear curve fitting and IC50 calculation was done with GraphPad Prism 9 using the inhibitor response three variable model or the exponential decay model. Images were prepared with Adobe Photoshop and assembled in Adobe Illustrator. All experiments presented were repeated at least twice independently. Data processing and reporting are adherent to the community standards.

#### SARS-CoV-2 PP assay

HEK293T-ACE2-GFP cells seeded in white, solid bottom 384-well microplates (Greiner BioOne) at 6,000 cells/well in 15 µL medium were incubated at 37°C with 5% CO_2_ overnight (∼16 h). Compounds were titrated 1:3 with 11 points in DMSO and dispensed into the assay plate at 23 nl/well via pintool. Cells were incubated with compounds for 1h at 37°C with 5% CO2 before 15 µl/well of PPs were added. The plates were then spinoculated by centrifugation at 1,500 rpm (453 x g) for 45 min and incubated for 48h at 37°C 5% CO2 to allow cell entry of PPs and the expression of luciferase. After the incubation, the supernatant was removed with gentle centrifugation using a Blue Washer (BlueCat Bio). Then 20 µL/well of Bright-Glo luciferase detection reagent (Promega) was added to assay plates and incubated for 5 min at room temperature. The luminescence signal was measured using a PHERAStar plate reader (BMG Labtech). Data were normalized with wells containing PPs as 100% and wells containing control DEnv PP as 0%.

#### ATP content cytotoxicity assay

HEK293T-ACE2-GFP cells were seeded in white, solid bottom 384-well microplates (Greiner BioOne) at 6,000 cells/well in 15 µl medium and incubated at 37°C with 5% CO_2_ overnight (∼16 h). Compounds were titrated 1:3 in DMSO and dispensed via pintool at 23 nl/well to assay plates. Cells were incubated for 1 h at 37°C 5% CO2 before 15 µl/well of media was added. The plates were then incubated at 37°C for 48h at 37°C 5% CO2. After incubation, 30 µl/well of ATPLite (PerkinElmer) was added to assay plates and incubated for 15 min at room temperature. The luminescence signal was measured using a Viewlux plate reader (PerkinElmer). Data were normalized with wells containing cells as 100%, and wells containing media only as 0%.

## Results and discussion

### The overall performance of the GCN model

Unlike traditional computational drug discovery methods such as structural homology-based drug search, the GCN classification model utilizes molecular graphs to extract spatial information. The modeling process computes in bonding environment at atomic or inter-atomic level within a fully connected framework as opposed to utilizing simple descriptors. As a result, the structural features of drug molecules can be well captured and built from low-level logic,^35,40^ making no emission of important possibilities. This method results in a robust performance with the classification accuracy as high as 99.5% for training set (the workflow was described in Figure 2). Additionally, the identified new compounds generally show structural dissimilarity to the training compounds, further highlighting its unique architecture compared to other structural assignment-based approaches.

**Figure 2:**
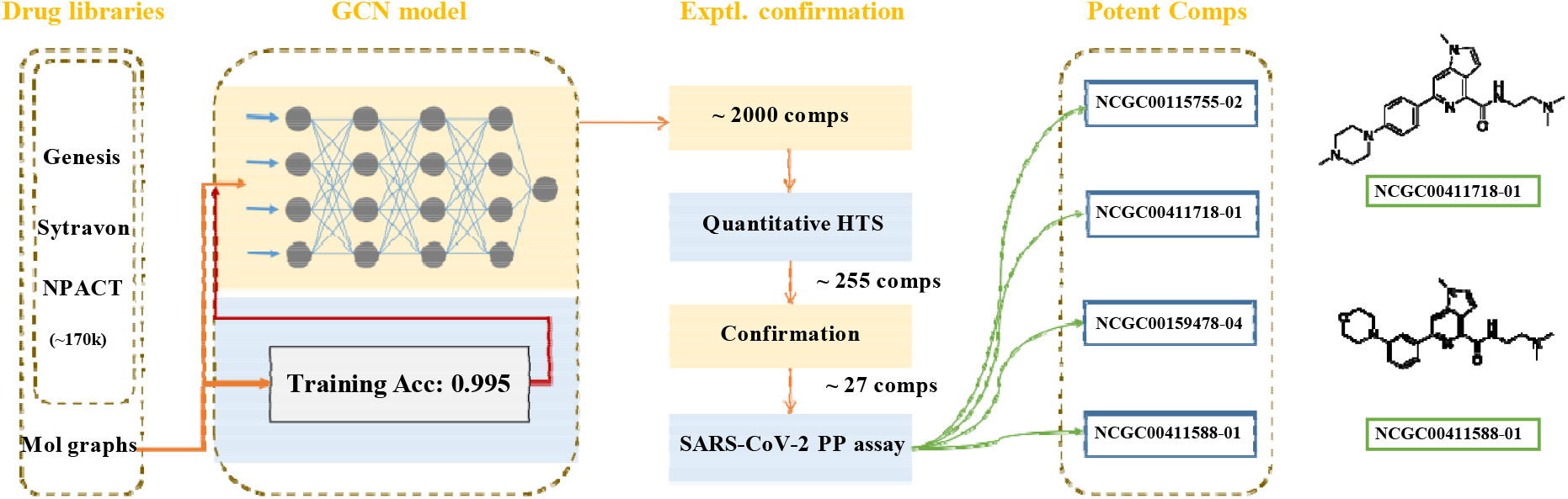
The workflow of GCN classification model upon endocytosis screenings.

### Identification of inhibitors for HSPG-mediated endocytosis

We used the GCN-based model to screen 170,000 compounds. ∼2000 compounds were short-listed by the virtual screen, which generated a small library that could be rapidly processed by a conventional quantitative high-throughput screen (qHTS) (Figure 3a). We then employed pHrodo red labeled *α*-Synuclein fibrils as an HSPG cargo in a combination screen because *α*-Synuclein fibrils share a similar entry mechanism as SARS-CoV-2.^13^ Importantly, the fluorescence intensity of cells treated with pHrodo-labeled *α*-Synuclein fibrils is only dependent on the amount of internalized cargo and the endolysosomal pH. By comparison, the luciferase-based pseudoviral entry assay can be influenced not only by the level of viral entry, but also by other factors that impact mRNA expression, translation, and luciferase stability. The screen identified 256 active compounds with most potent IC50 value of 0.95 uM. We cherry-picked 10 top compounds based on their potency and structural novelty (Figure 3b), and measured their cytotoxicity by an ATP content assay. The results showed that for 4 out of the 10 compounds, the IC50 for cytotoxicity was at least 10-fold larger than that for the inhibition of *a*-Synuclein fibril uptake (Figure 3b and c), suggesting a safety window for the usage of these drugs as endocytosis inhibitors.

**Figure 3:**
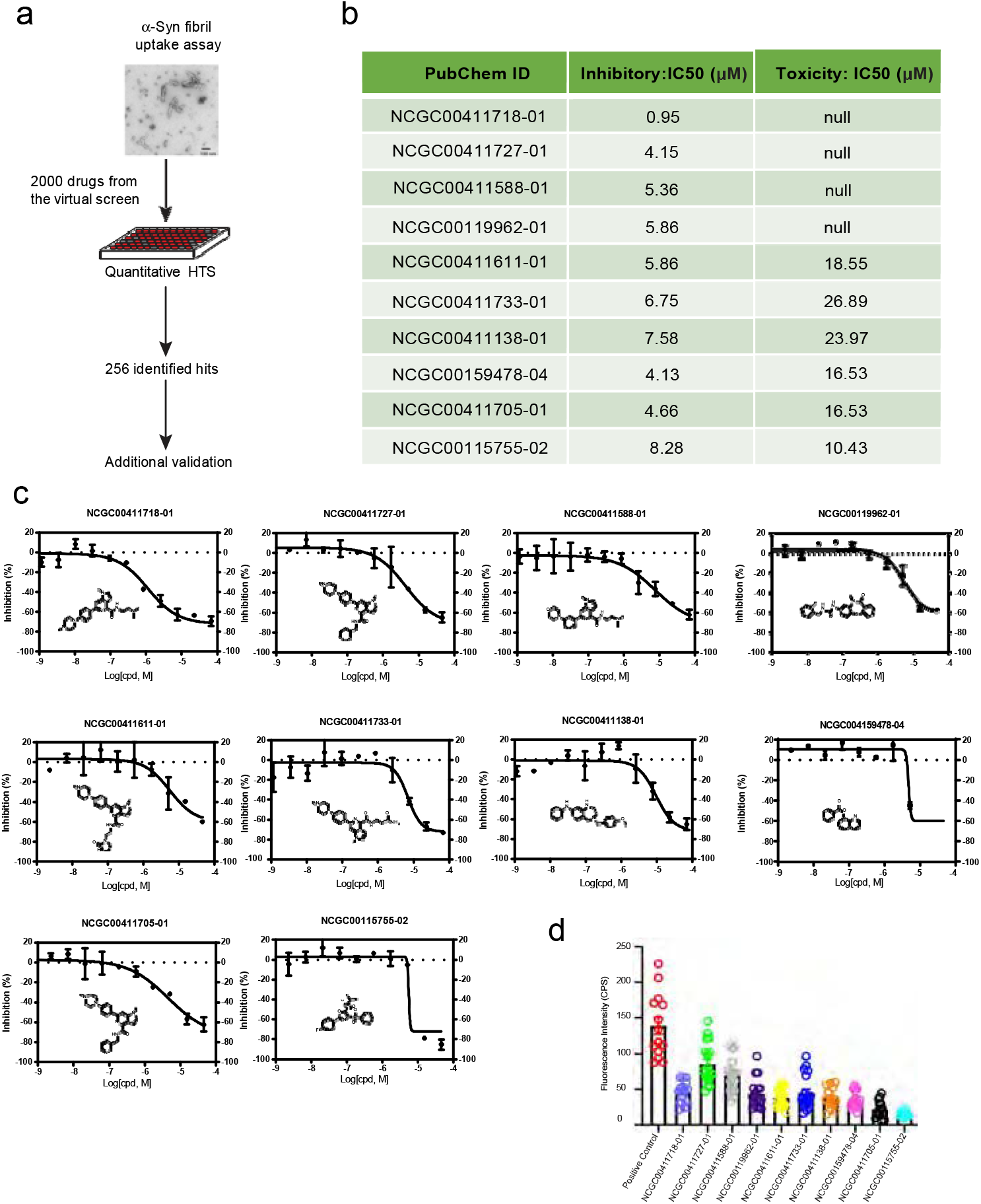
Identification of inhibitors for HSPG-mediated endocytosis: (a). The workflow of *α*-Synuclein fibrils uptake assay for confirmation of hits from virtual compound screen. (b). A summary of the activities of the top 10 compounds, IC50 was determined by titration experiments. (c). Dose-response curves of compound’s inhibitory effect on *α*-Synuclein fibrils uptake. (d). Measured fluorescence intensity of internalized *α*-Synuclein fibril-Alexa_596_ by U2OS cells treated with compound at 2-fold of its IC50. The experimental repeat number is 3.

To rule out false-positive hits due to compound-induced changes in lysosomal pH, which could reduce the fluorescence of internalized *α*-Synuclein fibrils, we measured the uptakes of *α*-Synuclein fibrils labeled with a pH-insensitive dye (Alexa_596_) in U2OS cells. When cells were treated with the top 10 inhibitors at concentrations 2-fold higher than their respective IC50 values, we found that all compounds tested could significantly inhibit the uptake of *a*-Synuclein fibrils compared to control treated cells (Figure 3d). These results suggest that these chemicals are indeed endocytosis inhibitors that block HSPG-mediated entry of *a*-Synuclein fibrils. We then treated cells with increased concentrations of NCGC00411718 and NCGC00159478, which showed the highest inhibition on the entry of pHrodo-labeled *a*-Synuclein fibrils. Drug-treated cells were incubated with Alexa_596_-labeled *α*-Synuclein fibrils in the presence of the inhibitor for 2 hours and imaged by a confocal microscope. The results suggest that both compounds inhibit *α*-Synuclein fibril uptake in a dose dependent manner with IC50 comparable to that measured by pHrodo-labeled *α*-Synuclein fibrils (Figure 4a-d).

**Figure 4:**
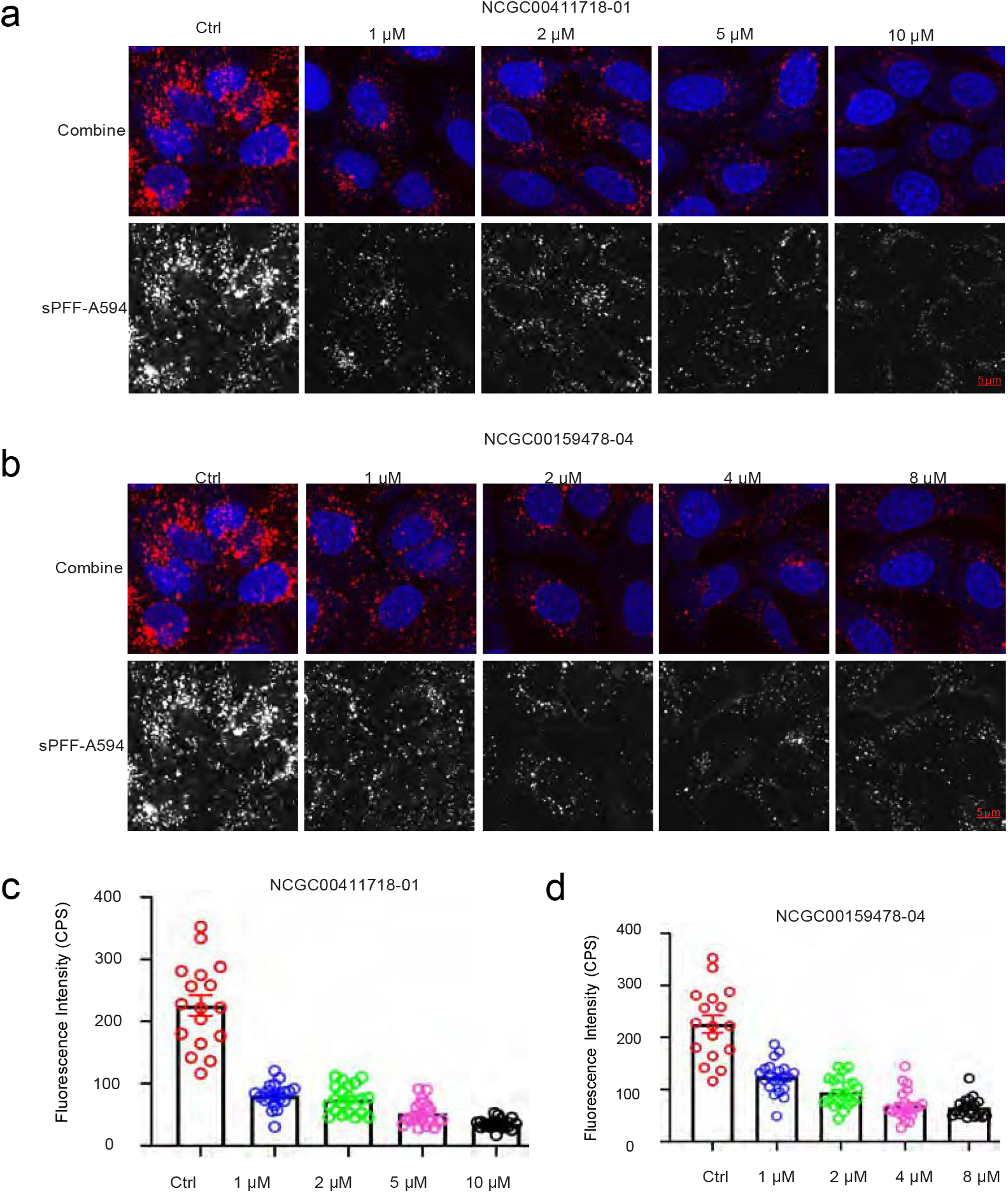
Identification of new endocytosis inhibitors targeting HSPG-mediated endocytosis. (a). NCGC00411718-01 inhibits of *a*-Synuclein fibril-Alexa594 uptake by U2OS cells in a dose dependent manner. (b). NCGC00159478-04 inhibits of *a*-Synuclein fibril-Alexa594 uptake by U2OS cells in a dose dependent manner. (c and d). Quantification of internalized *a*-Synuclein fibril-Alexa594 fluorescence intensity with compound treatment. Error bars indicate SEM. The experimental repeat number is 2.

### Identification of SARS-CoV-2 entry inhibitors

To test whether the newly identified endocytosis inhibitors could inhibit the entry of SARS-CoV-2, we used a previously established pseudotyped particle entry assay (Figure 5a). As shown previously,^6^ the entry of the pseudoviral particles into cells results in the expression of the luciferase reporter. To control the impact of ACE2-GFP expression levels on viral entry under drug-treated conditions, we normalized the luciferase signals by the ACE2-GFP level. We also measured the cytotoxicity of these chemicals in ACE2-GFP expressing cells using an ATP-based cell viability assay. We analyzed the top 27 compounds from the 256 inhibitors identified from the *α*-Synuclein fibril uptake screen. Among them, 16 in total showed an inhibitory activity against the viral entry with the most potent IC50 value of 0.76 µM. It is notable that some toxicity was observed for these compounds in HEK293T-ACE2-GFP cells after 48 hr treatment. The viral inhibition and cytotoxicity curves of the top 6 compounds are shown in Figure 5b.

**Figure 5:**
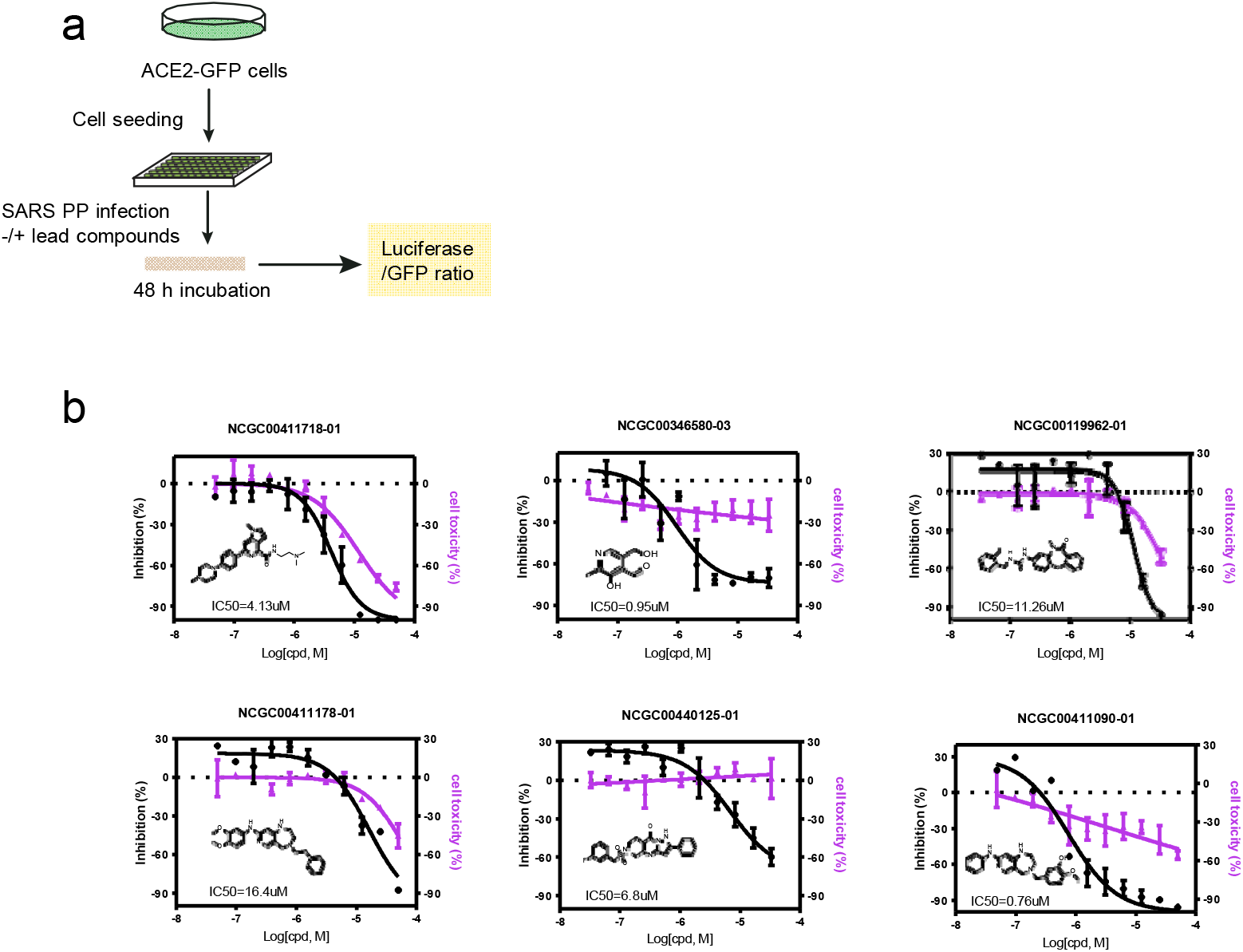
Identification of SARS-CoV-2 entry inhibitors: (a). The experimental scheme for inhibitor testing in HEK293T-ACE2-GFP cells. (b). Dose-responsive titration of compound’s inhibitory effect on SARS-CoV-2 entry and cytotoxicity. The experimental repeat number is 3.

### NCGC00115755 inhibited SARS-CoV-2 pseudotyped particle entry by disrupting actin filaments

We previously showed that the actin network under the plasma membrane is critical for the entry of HSPG-dependent endocytosis cargos including SARS-CoV-2. ^6,13^ We therefore asked whether any of the newly identified endocytosis could inhibit the actin cytoskeleton. To this end, we stained U2OS cells with Alexa488-labeled phalloidin, an actin binding dye. In control-treated cells, actin filaments were readily detected, which often run in parallel (Figure 6a). When cells treated with the top 10 endocytosis inhibitors were stained by Alexa488-labeled phalloidin, we observed dose-dependent disruption of cortex actin filaments only in NCGC00115755-02-treated cells by confocal fluorescence microscopy (Figure 6a) and it has anti-pseudotyped particle activity at IC50 of 5 M. Live cell imaging of cells expressing GFP-tagged Tractin, an actin binding reporter showed that untreated cells contain, in addition to stress fibers, many actin nucleation sites near the plasma membrane, which assemble comet tails (Supplementary videos). By contrast, in drug treated cells, the number of actin stress fibers were significantly reduced and actin comet tails were barely detectable (Supplementary videos). Altogether, these findings suggest that NCGC00115755-02 disrupts actin filament assembly, resulting in an endocytosis defect.

**Figure 6:**
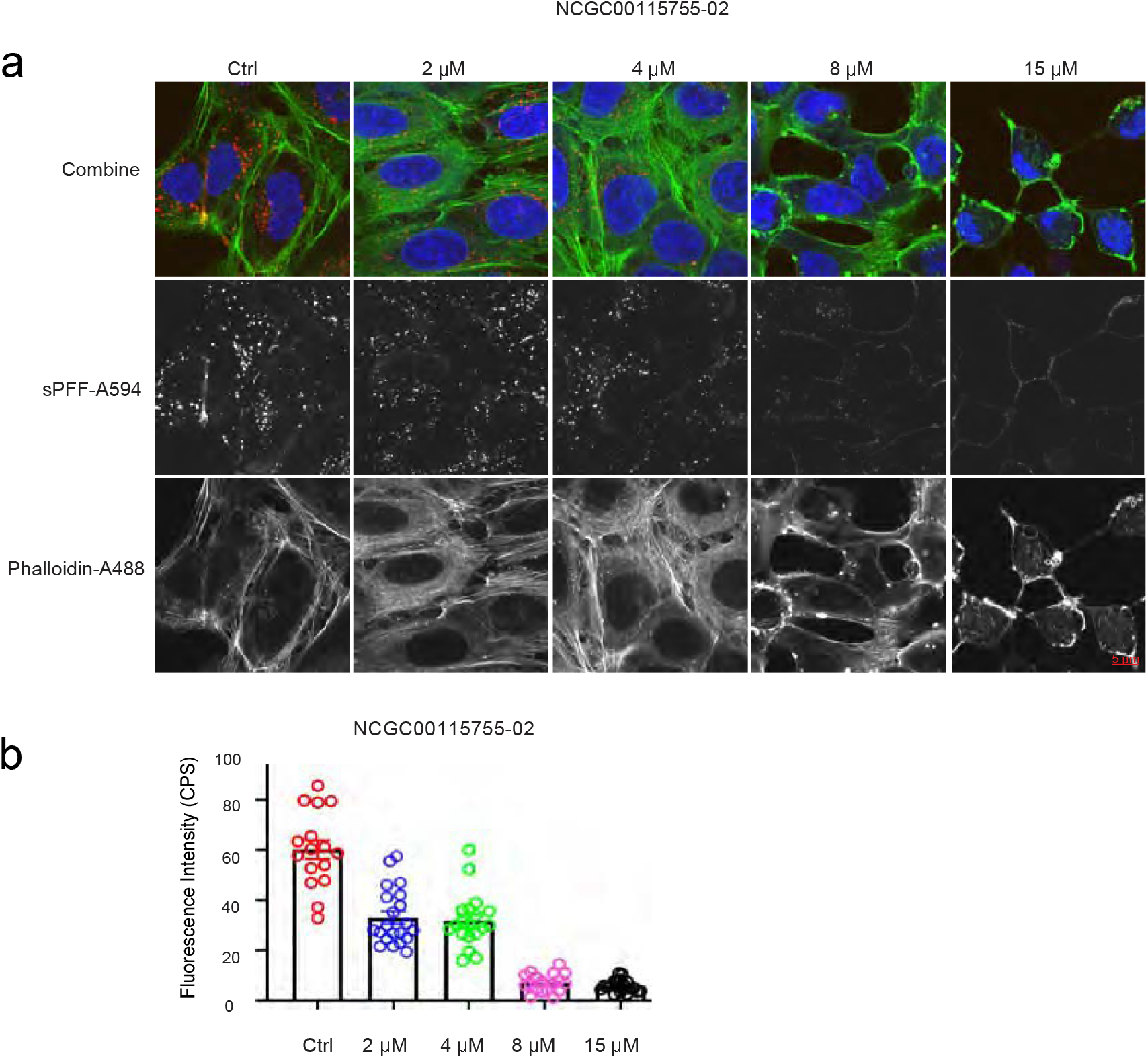
NCGC00115755-02 targets cellular actin cytoskeleton: (a). Cells treated with NCGC00115755-02 at the indicated concentrations were incubated with Alexa594-labeled *a*-Synuclein fibrils for 2 hours. Cells were stained with Phalloidin-Alexa488 in green to detect actin filaments and DAPI in blue to reveal the nuclei and then imaged. Note that cells treated with the drug has reduced level of internalized *a*-Synuclein fibrils. NCGC00115755-02 treatment also causes the disassembly of actin stress fiber and generates large actin aggregates. (b). Quantification of Alexa594-labeled *a*-Synuclein fluorescence intensity in a. Error bars indicate SEM. The experimental repeat number is 2.

## Conclusion

Machine learning-based virtual screening technologies have the potential to efficiently select drug candidates for specific targets with high accuracy at an affordable cost, and therefore, is an important complementary strategy to conventional high-throughput small molecule screening (HTS). SARS-CoV-2 viruses co-opt a cellular endocytosis pathway to enter human airway epithelial cells. This key viral entry step has been subjected to conventional drug repurposing screens, yielding several viral entry inhibitors. In this study, we developed and trained a GCN model using the structural information from previously identified SARS-CoV-2 entry inhibitors. When this model was applied to untested chemical libraries, it can efficiently select compounds with high probability of showing an anti-SARS-CoV-2 activity. This model, when combined with conventional drug screening assays, generates a powerful platform that allows rapid identification of new SARS-CoV-2 entry inhibitors. In principle, this platform can be applied to any drug targets, which can quickly expand the existing inhibitor repertoire of any class. The findings shown in this study have revealed a promising venue for accelerated drug development.

## Acknowledgement

The work was supported by the intramural research program of the National Institute of Diabetes, Digestive & Kidney Diseases (Y.Y.) and by the National Center for Advancing Translational Sciences (W.Z.) in the National Institutes of health.

## Data and software availability

Technical details of the developed package can be found on our GitHub page: github.com/tcsnfrank0177/Graph-convolutional-network-DrugScreening.git. Programming environment: Python 3.6 or higher is recommended. Supplementary videos are provided as attachment.

## Notes

### Competing Interest Statement

The authors have declared no competing interest.

